# multiomeR: a pipeline for reproducible analysis of large-scale single-nucleus Epi Multiome datasets

**DOI:** 10.64898/2026.10.14.759117

**Authors:** Thomas Gade Koefoed, Shanshan He, Malte Thodberg, Laura Maarit Pikkupeura, Kata Krizic, Benedicte Schultz Kapel, Raquel Sanz Martinez, Simon Rasmussen, Torben Hansen

## Abstract

**Summary:** The 10x Genomics Epi Multiome assay measures gene expression and chromatin accessibility in the same nuclei, and studies increasingly profile many samples. Yet analysis after Cell Ranger ARC is usually assembled by hand from separate tools, which makes it slow to revise, hard to reproduce and memory-intensive for large studies. multiomeR addresses these problems with a configurable R pipeline for quality control, processing and integration of both data types, and cell-type annotation across many samples. Plain-text configuration files record every analytical choice, each step is tracked so that a change reruns only what it affects, and data stay on disk rather than in memory. Optional modules compare groups of donors, link accessible regions to the genes they may regulate, and test which cell types are enriched for genetic associations with a trait. In a benchmark on 31 public samples (375,757 nuclei), multiomeR had a shorter estimated run time than a Seurat v5.6/Signac v2 workflow (2.2 versus 5.6 h), because it runs independent steps in parallel. It also used fewer memory gigabyte-hours (87 versus 114 GB-hours), with similar total CPU time. multiomeR thus makes large multi-sample multiome studies practical to analyze, revise and extend.

**Availability and implementation:** multiomeR is implemented in R and runs on Linux (x86-64). Pixi installs fixed versions of all required software. It is freely available under the MIT license at https://github.com/koefoeden/multiomeR. The version described here, 1.2.2, is archived at https://doi.org/10.5281/zenodo.23369832, and the user manual is at https://koefoeden.github.io/multiomeR/.

**Supplementary information:** Supplementary data are available online.

## 1 Introduction

The 10x Genomics Epi Multiome assay measures gene expression and chromatin accessibility (ATAC, assay for transposase-accessible chromatin) in the same nuclei, so regulatory DNA activity can be linked to transcription. Seurat, Signac, ArchR, SnapATAC2 and muon provide methods to analyze such data (Granja et al. 2021, Hao et al. 2021, Stuart et al. 2021, Bredikhin et al. 2022, Hao et al. 2024, Zhang *et* al. 2024). Some of them store data on disk or organize several samples in one project, but none provides a ready-made workflow from Cell Ranger ARC (10x Genomics 2026) outputs of many samples to donor-level results (Supplementary Table S1). Researchers therefore write and maintain their own scripts to connect quality control, clustering, cell-type annotation and downstream analyses. Such scripts take time to write, are prone to errors and are hard to reproduce.

multiomeR replaces these scripts with a configurable pipeline from Cell Ranger ARC outputs to donor-level results. Plain-text configuration files with defaults record every analytical choice. Every step is tracked with targets (Landau 2021), so a change reruns only the steps it affects, and review checkpoints let users inspect results before continuing. Data stay on disk, which limits memory use as studies grow, and the downstream analyses treat donors, not nuclei, as replicates.

## 2 Overview of the pipeline

multiomeR has a primary module and three optional downstream modules (Supplementary Figure S1). The downstream modules cover the analyses that multi-donor Epi Multiome studies most often perform after processing: comparing groups of donors, linking accessible regions to genes, and relating cell types to genetic associations. multiomeR supports human and mouse data. The genetic enrichment module requires human data, and the donor-level analyses require several donors, the individuals that the samples come from, per group or cell type.

### 2.1 Primary module

Samples from different tissues and protocols differ in quality, cell composition and ambient RNA, so settings such as quality-control thresholds and clustering resolutions are best chosen after looking at the data. The primary module therefore takes the Cell Ranger ARC outputs of one or more samples to annotated cell types in eight stages, each ending at a review checkpoint (Supplementary Figure S1). Each stage asks only for the settings it needs and ends with plots whose captions say what to look for. Users can then accept the result, or revise those settings and rerun the stage, before moving on, and defaults cover everything else. Along the way, the module can assign nuclei of pooled samples to donors by genotype (Huang Y et al. 2019, Huang X and Huang 2021) and use gene-expression counts corrected for ambient RNA with CellBender (Fleming et al. 2023). It integrates both data types by weighted nearest-neighbor (WNN) integration (Hao *et al*. 2021) and annotates cell types from marker genes. Supplementary Methods describes each step, Supplementary Table S5 lists the tools used, and the user manual lists every setting.

### 2.2 Differential analyses module

Multi-donor studies are usually designed to compare groups of donors, such as cases and controls, often while adjusting for covariates. The differential analyses module defines each comparison as a short configuration entry: an R model formula, its contrasts and, optionally, the cell types or samples to include. One configuration can therefore hold several study designs. For each comparison, the module tests for changes in cell-type composition (Brooks et al. 2017). Within each cell type, it tests gene expression, chromatin accessibility, motif accessibility and inferred transcription factor activity (Schep et al. 2017, Badia-i-Mompel et al. 2022, Müller-Dott et al. 2023) with established pseudobulk methods that treat donors as replicates (Robinson et al. 2010, Ritchie et al. 2015). Gene-set tests summarize the gene-expression results (Wu and Smyth 2012, Liberzon *et al*. 2015, Milacic *et al*. 2024, Dolgalev 2026). Supplementary Methods describes the models.

### 2.3 Peak–gene correlation module

Because both data types are measured in the same nuclei, Epi Multiome data can link candidate regulatory regions to the genes they may control, a key step in interpreting non-coding genetic variants. The peak– gene correlation module tests, within each cell type, whether the accessibility of a peak and the expression of a nearby gene vary together across donors and cell states. It treats donors rather than nuclei as replicates, so that thousands of nuclei from a few donors are not counted as independent evidence. Supplementary Methods describes the model and its calibration.

### 2.4 Genetic enrichment module

Most variants associated with complex traits lie in non-coding regions that act in specific cell types. The cell types in which these variants are accessible can therefore point to the cells through which they act. The genetic enrichment module takes fine-mapped results of genome-wide association studies (GWAS) from the Open Targets Platform, or the user’s own (Buniello et al. 2025, Cerezo et al. 2025). It tests whether peaks containing likely causal variants are more accessible in a cell type than comparable background peaks (Schep et al. 2017, Ulirsch et al. 2019, Germain 2026). It breaks each enrichment down into its contributing loci and variants, showing whether it rests on many loci or a few. A reimplementation of SCAVENGE extends the analysis to individual nuclei (Yu et al. 2022). Supplementary Methods describes the tests.

### 2.5 Outputs

Results are only useful if they can be inspected, reused and published. multiomeR therefore saves intermediate and final results, exports Seurat/Signac objects for use with other tools, and keeps the data behind every quality-control and result plot, so the plots can be redrawn for publication.

## 3 Example application

To illustrate the outputs, we analyzed 31 public human Epi Multiome samples from 10x Genomics and ENCODE, covering six tissues and blood (Supplementary Table S2) (ENCODE Project Consortium *et al*. 2020, Luo et al. 2020). The primary module kept 213,212 of 375,757 nuclei (57%) after quality control and assigned them to 19 cell types or left them unassigned. Marker genes distinguish these cell types in both gene expression and promoter accessibility (Figure 1A). The peak–gene correlation module’s link with the lowest P-value connects a peak and the *SGPL1* gene in NK cells from 11 donors (Figure 1B–C). The genetic enrichment module tested 26 GWAS with enough fine-mapped signal in peaks (Supplementary Table S3). Of these, 12 were enriched in at least one cell type (Benjamini–Hochberg q < 0.05), involving 10 of the 19 cell types (Supplementary Figure S3). Figure 1D shows 8 of these studies, such as low-density lipoprotein (LDL) cholesterol (Graham et al. 2021), enriched in hepatocytes, and colorectal cancer (Law et al. 2019), enriched in colonocytes and goblet cells. Both rest on many loci. The LDL signal is spread over 360 loci with variants in peaks, and the largest of 51 such colorectal cancer loci contributes 32% of its signal (Figure 1E). For colorectal cancer, the module’s scores for individual nuclei are highest in goblet cells and colonocytes (Figure 1F). Supplementary Methods lists the settings used.

**Figure 1:**
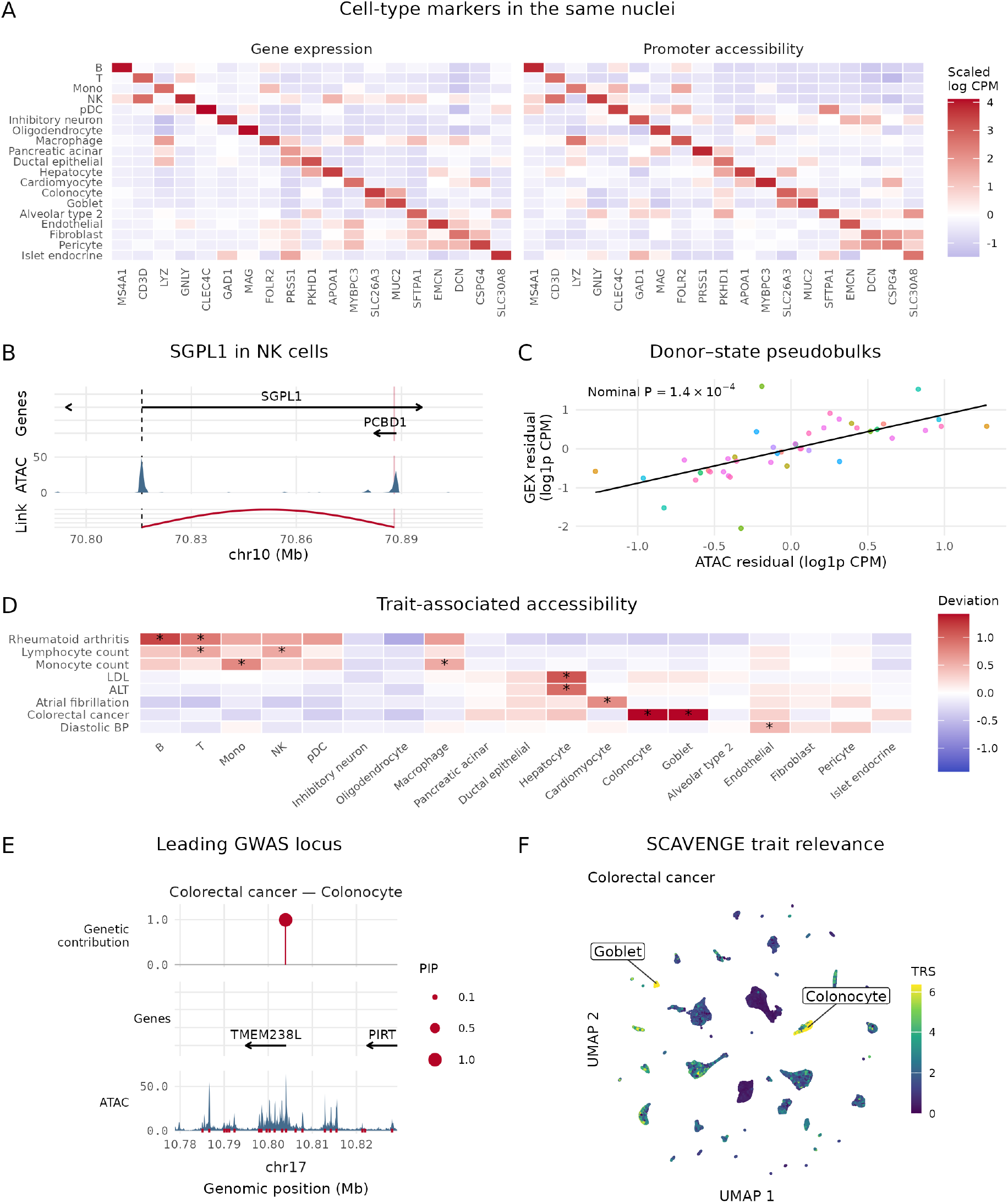
Selected multiomeR outputs from 31 public human Epi Multiome samples. **(A)** Expression and promoter accessibility of a marker gene of each cell type, in cell-type pseudobulks summed over donors, as log counts per million (CPM) scaled across cell types. **(B)** The peak–gene link with the lowest P-value: genes, ATAC coverage in NK cells, the linked peak (red) and the *SGPL1* transcription start site (dashed). **(C)** Expression against accessibility of donor–state pseudobulks, adjusted for donor, sequencing depth and the cell type’s main chromatin states. Points are colored by donor, with the nominal P-value of the hierarchical model and a line fitted for illustration. **(D)** Accessibility of trait-associated peaks in each cell type relative to background peaks (chromVAR deviation, Schep et al. 2017), for 8 GWAS traits whose enrichment rests on many loci (Law et al. 2019, Vuckovic et al. 2020, Graham et al. 2021, Ishigaki et al. 2022, Ghouse et al. 2024, Keaton et al. 2024, Roselli et al. 2025). Stars mark Benjamini–Hochberg q < 0.05. Supplementary Figure S3 shows all tested traits. **(E)** The locus contributing most to the colorectal cancer–colonocyte association: each variant’s contribution (vertical line) and fine-mapping posterior inclusion probability (PIP; point size), protein-coding genes, ATAC coverage and peaks (red). **(F)** SCAVENGE trait-relevance scores (TRS) of colorectal cancer for individual nuclei, shown on a uniform manifold approximation and projection (UMAP) (McInnes et al. 2018) of the WNN graph and labeled by the cell types enriched for it. ALT, alanine aminotransferase; BP, blood pressure; GEX, gene expression; LDL, low-density lipoprotein; Mono, monocytes; NK, natural killer cells; pDC, plasmacytoid dendritic cells.

## 4. Efficiency and scaling

Because studies grow to hundreds of thousands of nuclei, multiomeR keeps count matrices and ATAC fragments on disk with BPCells (Parks and Greenleaf 2025). It runs the analysis as many small steps, in parallel on a workstation or a computing cluster (Landau 2021, 2026). We also reimplemented four established methods to use less memory: UCell, AMULET, WNN integration and SCAVENGE (Andreatta and Carmona 2021, Hao et al. 2021, Thibodeau et al. 2021, Yu et al. 2022). They reproduce the original results exactly or closely (Supplementary Table S4). Our AMULET implementation, for example, matched scDblFinder’s exactly while running 71 times faster with 97% less peak memory (Germain et al. 2022). multiomeR runs on Linux with at least 32 GB of memory and fixed software versions from Pixi (Fischer et al. 2025), and its two-sample demo runs in about 30 minutes with 16 threads.

We compared resource use with a Seurat v5.6/Signac v2 workflow that also keeps data on disk (Supplementary Figure S2; Supplementary Methods). Both workflows took 1 to 31 example samples from Cell Ranger ARC outputs to joint clustering and embedding, only part of the primary module. For 31 samples (375,757 nuclei), multiomeR’s critical path, its run time with independent steps in parallel, was 2.2 h, against 5.6 h for Seurat/Signac. Both used similar CPU time (11.3 versus 11.3 CPU-hours) and peak memory, whereas multiomeR used fewer memory gigabyte-hours (87 versus 114 GB-hours) and more disk space, mainly for a BPCells copy of the ATAC fragments (Supplementary Figure S2).

## 5. Conclusion

Epi Multiome studies are moving from a few samples to many donors, and their analysis must become as scalable and reproducible as their data generation. multiomeR provides one configurable workflow from Cell Ranger ARC outputs to cell types, donor-level comparisons, candidate enhancer–gene links and trait-relevant cell types. Every analytical choice is recorded, steps are rerun only when a change requires it, and data stay on disk. This lets researchers spend their time interpreting multiomic data rather than maintaining analysis scripts.

## Code and data availability

multiomeR is available under the MIT license at https://github.com/koefoeden/multiomeR, with a user manual at https://koefoeden.github.io/multiomeR/. The version described here, 1.2.2, is archived at https://doi.org/10.5281/zenodo.23369832. The public Epi Multiome datasets are available from 10x Genomics and the ENCODE portal under the identifiers in Supplementary Table S2. The GWAS credible sets come from the Open Targets Platform, release 26.03 (Supplementary Table S3). The archived release also contains the configuration (configuration_dev/) and code (pipeline_manuscript_figures/) behind every figure and table of results, together with their source data and the values that the text quotes.

## Acknowledgements

HPC facilities and technical expertise were provided by the Data Analytics Platform at the Novo Nordisk Foundation Center for Basic Metabolic Research. Coding assistants based on large language models helped write and review code and documentation. For the manuscript, they were used to improve clarity for readers and to check the text against the code and results; the authors determined its content and conclusions, reviewed all output and take full responsibility for the manuscript.

## Author contributions

T.G.K.: Conceptualization (equal), Data curation (lead), Formal analysis (lead), Funding acquisition (lead), Investigation (lead), Methodology (lead), Project administration (equal), Software (lead), Validation (equal), Visualization (lead), Writing – original draft (lead), Writing – review & editing (lead).

S.H.: Data curation (supporting), Investigation (supporting), Methodology (supporting), Software (supporting), Validation (equal), Visualization (supporting), Writing – review & editing (supporting).

M.T.: Conceptualization (equal), Investigation (supporting), Methodology (supporting), Software (supporting), Project administration (equal), Supervision (equal), Validation (equal), Writing – review & editing (supporting).

L.M.P.: Conceptualization (equal), Investigation (supporting), Methodology (supporting), Software (supporting), Project administration (equal), Supervision (supporting), Validation (equal), Writing – review & editing (supporting).

K.K.: Conceptualization (supporting), Writing – review & editing (supporting). B.S.K.: Conceptualization (supporting), Writing – review & editing (supporting).

R.S.M.: Data curation (supporting), Investigation (supporting), Methodology (supporting), Software (supporting), Validation (equal), Visualization (supporting), Writing – review & editing (supporting).

S.R.: Supervision (equal), Writing – review & editing (supporting).

T.H.: Funding acquisition (supporting), Resources (lead), Supervision (equal), Writing – review & editing (supporting).

## Supplementary material

Supplementary Methods, Supplementary Figures S1–S3 and Supplementary Tables S1–S6 are available online.

