## Supplementary Figures S1-S3 for "multiomeR: a pipeline for reproducible analysis of large-scale single-nucleus Epi Multiome datasets"

### multiomeR: a pipeline for reproducible analysis of large-scale single-nucleus Epi Multiome datasets

Thomas Gade Koefoed[1,*], Shanshan He[1], Malte Thodberg[1], Laura Maarit Pikkupeura[1], Kata Krizic[1], Benedicte Schultz Kapel[1], Raquel Sanz Martinez[1], Simon Rasmussen[1], Torben Hansen[1,*]

[1] Novo Nordisk Foundation Center for Basic Metabolic Research, Faculty of Health and Medical Sciences, University of Copenhagen, Blegdamsvej 3B, 2200 Copenhagen N, Denmark

**This file includes:**

Supplementary Figure S1: Organization of the multiomeR workflow
Supplementary Figure S2: multiomeR and a Seurat/Signac baseline with data on disk use similar CPU time, but multiomeR has a shorter critical path and uses fewer memory gigabyte-hours
Supplementary Figure S3: Cell-type enrichment of all 26 GWAS in the example application

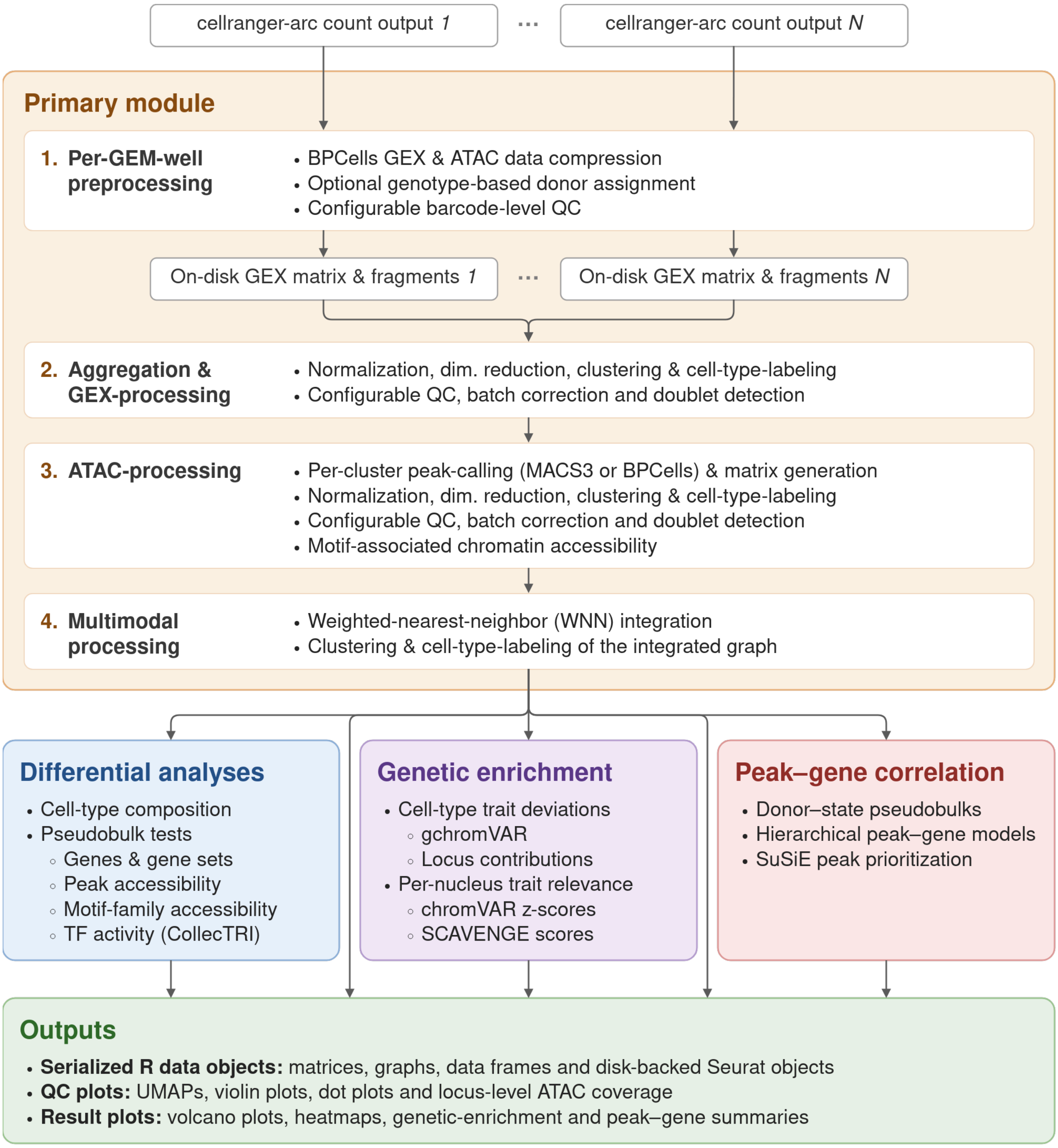


**Supplementary Figure S1. Organization of the multiomeR workflow.** The primary module (orange) takes the Cell Ranger ARC (10x Genomics 2026) outputs of each sample to an integrated, annotated aggregation of samples in four parts, which contain its eight review checkpoints. The three optional modules (blue, purple, red) start from its results, and all modules write the outputs listed in green. Supplementary Methods describes each step, and Supplementary Table S5 lists the tools.

**Abbreviations:** GEX, gene expression; ATAC, assay for transposase-accessible chromatin; QC, quality control; dim., dimensionality; TF, transcription factor; SuSiE, sum of single effects (Wang *et al.* 2020).

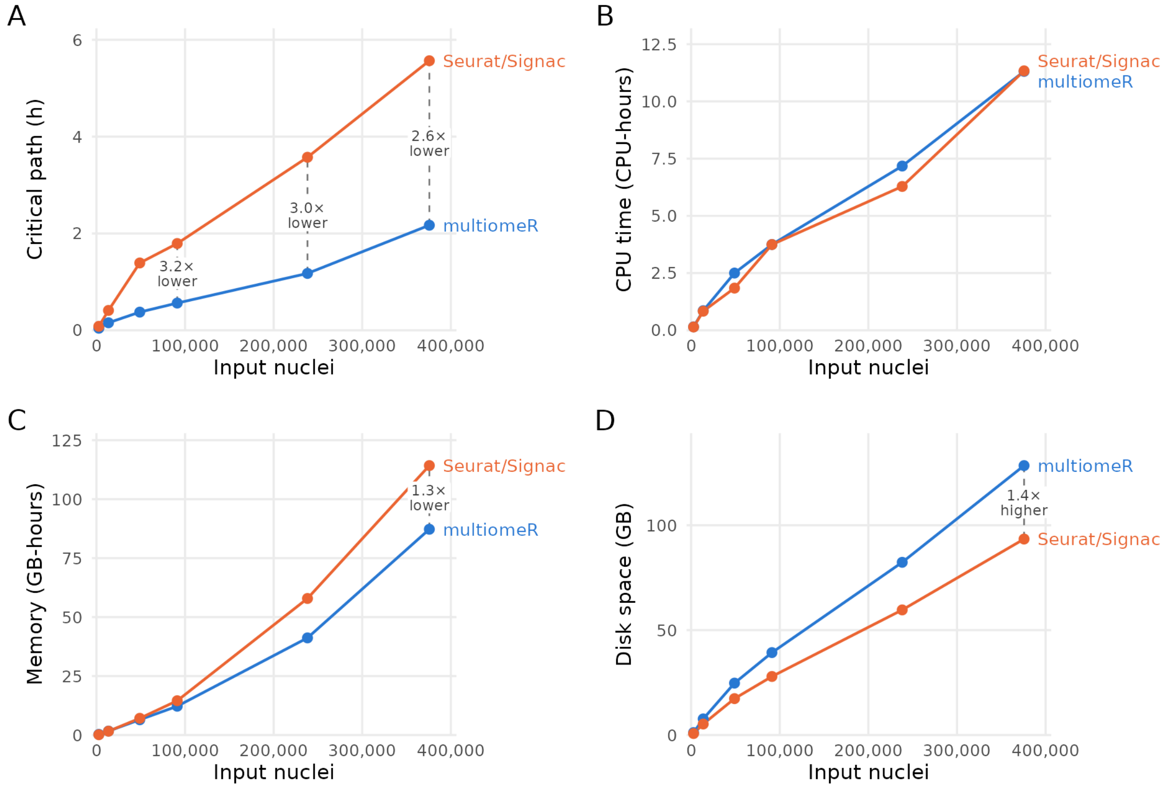


**Supplementary Figure S2. multiomeR and a Seurat/Signac baseline with data on disk use similar CPU time, but multiomeR has a shorter critical path and uses fewer memory gigabyte-hours.** Both workflows ran from Cell Ranger ARC outputs to joint WNN clustering and embedding in a multimodal Seurat/Signac object (Stuart *et al*. 2021, Hao *et al*. 2024), on nested aggregations of 1 to 31 samples of Supplementary Table S2 (2,713 to 375,757 nuclei). multiomeR's values exclude analyses that only it performs. **(A)** Critical path: the estimated run time with unlimited parallel jobs for multiomeR and with steps run in sequence for Seurat/Signac. **(B)** CPU time. **(C)** Memory gigabyte-hours. **(D)** Disk space, including the Cell Ranger outputs that both read. Dashed lines mark differences that span at least a fifth of a panel's height, labeled with the fold difference of multiomeR from Seurat/Signac. Each workflow ran once. Supplementary Methods describes the design, and Supplementary Table S6 lists every value.

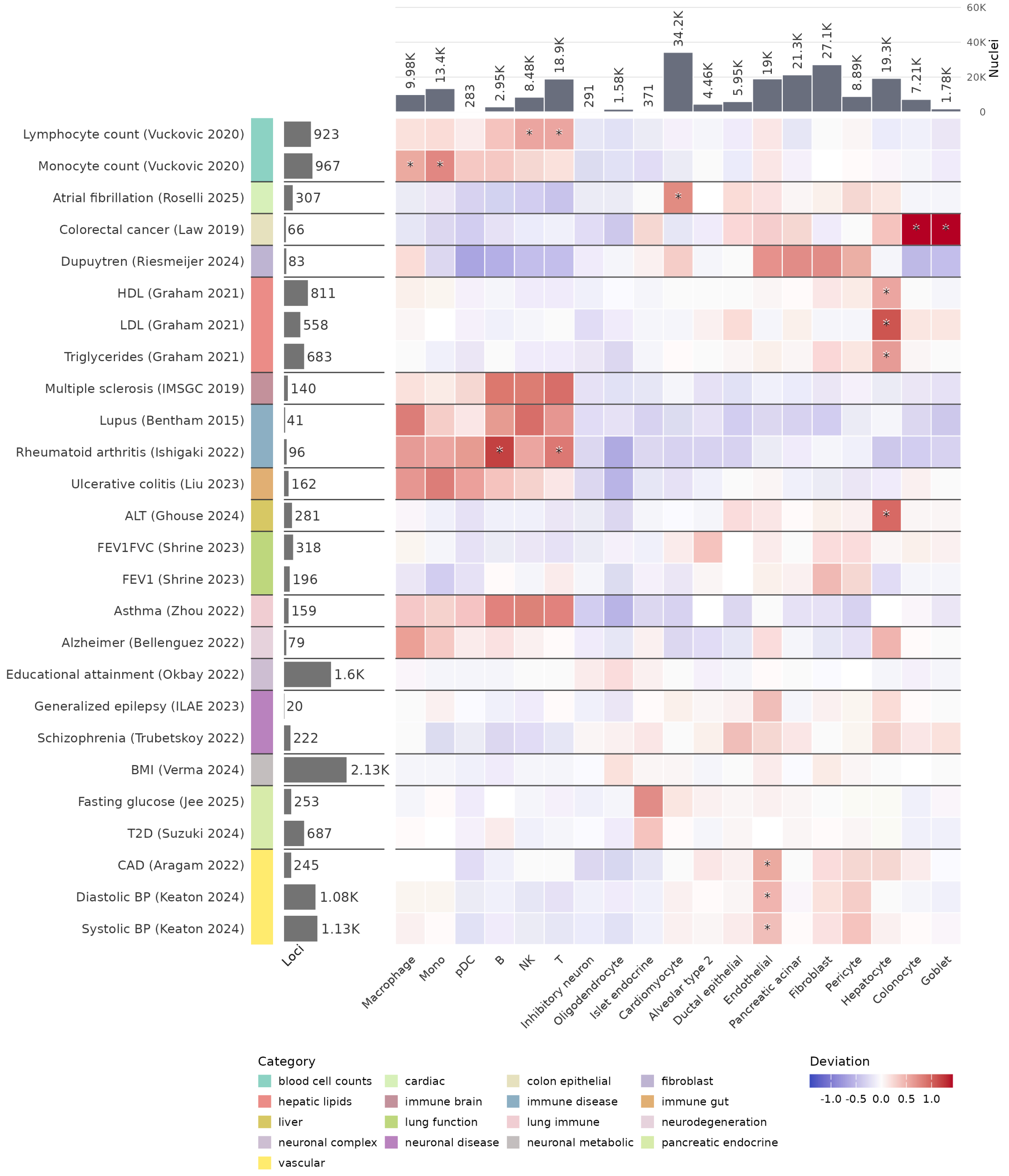


**Supplementary Figure S3. Cell-type enrichment of all 26 GWAS in the example application.** Rows are studies, grouped by trait category, with the trait category and the number of credible-set loci on the left. Supplementary Table S3 lists each study by the same trait, first author and year, with its fine-mapping method and sample size. Columns are the 19 assigned cell types, with their numbers of nuclei above. Colors show the chromVAR deviation (Schep *et al.* 2017) of each study's trait-associated peaks in each cell type, on the color scale of Figure 1D. Stars mark $q < 0.05$, as in Figure 1D.
