## Supplementary Methods for "multiomeR: a pipeline for reproducible analysis of large-scale single-nucleus Epi Multiome datasets"

These methods describe multiomeR version 1.2.2 (commit `ca5d8f2`), archived at https://doi.org/10.5281/zenodo.23369832.

#### Workflow organization and execution

multiomeR consists of a primary module and optional modules for differential analyses, peak–gene correlation and genetic enrichment (Supplementary Figure S1). Two linked configuration layers define the inputs. Each sample specifies its Cell Ranger ARC (10x Genomics 2026) count directory and quality-control exclusions. Each aggregation selects samples for joint analysis and links their donor and sample metadata. Each sample is one GEM well, a single Chromium channel, and the methods below use that term.

Processing follows standard practice for each data type (Heumos *et al.* 2023). It includes per-sample quality control, principal component analysis (PCA) of gene expression, latent semantic indexing (LSI) of accessibility, Leiden clustering (Traag *et al.* 2019) and separate unimodal analyses before weighted nearest-neighbor integration. The following sections describe each analysis step with its main fixed settings. They also form the methods pages of the user manual (https://koefoeden.github.io/multiomeR/). These pages link each step to its source files, and the manual's parameter browser lists every setting and its default. The primary module's eight review checkpoints (Supplementary Figure S1) let users inspect results and revise settings. Seurat/Signac objects (Stuart *et al.* 2021, Hao *et al.* 2024) exported after GEX processing and after multimodal integration allow analysis with other tools.

Each analysis step is a target in a dependency graph, so targets reruns only the steps whose code, inputs or settings have changed (Landau 2021). crew runs independent steps in parallel on a workstation or through a scheduler (Landau 2026). BPCells keeps large matrices and fragments on disk (Parks and Greenleaf 2025), and a locked Pixi environment fixes software versions (Fischer *et al.* 2025). Supplementary Table S5 lists the main tools.

multiomeR identifies each sample's reference from the hashes in its fragment file. It includes metadata for the human GRCh38 (2020-A and 2024-A) and mouse mm10 (2020-A) Cell Ranger ARC references. All samples of an aggregation must share one reference. It runs on Linux x86-64 with at least 32 GB of RAM, on a workstation or through a Slurm, PBS, SGE or LSF scheduler. Its most memory-intensive steps need about 7 GB plus 2.5 GB per 100,000 nuclei. The optional Seurat SCTransform backend (Hafemeister and Satija 2019, Choudhary and Satija 2022) for gene-expression normalization needs about 0.25 GB per 1,000 nuclei. The genetic enrichment module requires human GRCh38 data.

#### Preprocessing and sample quality control

Each GEM well supplies a Cell Ranger ARC count directory. Gene-expression counts are imported into BPCells from the filtered feature-barcode matrix, or from a CellBender output when configured (Fleming *et al.* 2023, Parks and Greenleaf 2025). ATAC fragments are imported from the Cell Ranger fragment file. Both carry a GEM-well-prefixed barcode, so nuclei remain distinct across wells. Per-well quality-control metrics are calculated for the barcodes that Cell Ranger called as cells. BPCells calculates ATAC metrics such as TSS enrichment and nucleosome signal. Gene-expression metrics, such as the fraction of counts from mitochondrial genes (names beginning with `MT-`, ignoring case), are calculated from the imported matrix. Optional genotype demultiplexing runs cellsnp-lite on the ATAC alignments, with a minimum minor-allele frequency of 0.1 and 20 reads per SNP (Huang X and Huang 2021). Vireo then demultiplexes the barcodes with the supplied donor genotypes (Huang Y *et al.* 2019). The most likely singlet donor replaces the GEM-well donor identifier of every barcode. Vireo's singlet, doublet and unassigned calls and probabilities are recorded as metadata for the exclusions. A BPCells-based implementation of AMULET reads the stored fragments of up to 1,000 bp one by one, without loading them all into memory (Thibodeau *et al.* 2021). This avoids the chromosome-scale `GRanges` objects that scDblFinder builds from the text fragment file. It skips fragments on chrM, chrX and chrY and, for GRCh38, fragments in the AMULET authors' list of repeats, segmental duplications and blacklisted regions, where multi-mapping reads cause spurious overlaps. It reproduces the overlap metrics and q-values of `scDblFinder::amulet()` but applies no automatic filter (Germain *et al.* 2022). Because AMULET performs well only on deeply sequenced nuclei, filtering on its q-values is left to the user, and its authors recommend $q < 0.01$ (Thibodeau *et al.* 2021).

Configured GEM-well exclusions are filter expressions applied to the called cells of each well. Demultiplexing and AMULET results can therefore be used as exclusion criteria before aggregation-level analysis. Called cells missing from the imported gene-expression matrix are always excluded. The remaining nuclei of the selected wells are combined without further filtering.

### GEX processing

#### Normalization and PCA

For each aggregation, the quality-controlled gene-expression matrices are combined with their GEM-well-prefixed barcodes. Genes with 50 or fewer total counts are excluded before dimensional reduction. Normalization and dimensional reduction use either the disk-backed BPCells Pearson-residual workflow or Seurat `SCTransform` (Hafemeister and Satija 2019, Parks and Greenleaf 2025). With the BPCells backend, analytic Pearson residuals are computed with one overdispersion parameter ($\theta = 100$) for all genes and clipped to $\pm\sqrt{N}$ for $N$ nuclei (Lause *et al.* 2021). Per-gene estimates of $\theta$ would absorb differences between cell types and flatten marker-gene residuals, and the $\sqrt{N}$ clip keeps markers of rare populations (Lause *et al.* 2021). Configured cell-level covariates can be regressed out. The 3,000 genes with the highest residual variance are kept, and a truncated singular-value decomposition gives the principal components. The Seurat backend applies SCTransform with its v2 regularization to the same cells and genes (Choudhary and Satija 2022). It keeps the same number of variable features and computes the principal components from the residual matrix. A cell-cycle difference score can be regressed out but is not by default. Its S and G2M scores are computed from log-normalized counts with the 2019 Seurat cell-cycle gene sets (Tirosh *et al.* 2016).

#### Batch correction and clustering

Harmony correction can be applied separately to the selected GEX principal components and ATAC LSI dimensions, with at most 25 iterations and `lambda = 1` (Korsunsky *et al.* 2019, Patikas *et al.* 2026). It is off by default, since correcting a variable can also remove biology confounded with it (Luecken *et al.* 2022). The configured metadata columns are combined into one interaction batch factor, and the ATAC correction can add its own columns. Nuclei with missing covariate values are removed with a warning.

For each modality, approximate nearest neighbors are found on the selected dimensions with BPCells hierarchical navigable small world (HNSW) search (Malkov and Yashunin 2020), using cosine distance. The neighbors are converted to a shared-nearest-neighbor (SNN) graph with Jaccard weights, removing edges with weights below 1/15. The graph is clustered with the modularity objective of the Leiden algorithm in ten iterations (Traag *et al.* 2019, Parks and Greenleaf 2025). Clusters are renumbered by size, and clusters of fewer than 100 nuclei by default are removed from the GEX, ATAC and WNN clusterings. UMAP embeddings are computed with uwot using cosine distance on separately configurable dimensions, by default the clustering dimensions (McInnes *et al.* 2018, Melville 2026).

#### Cell-type annotation

Cell-type annotation uses GEX marker signatures supplied by the user. Signatures list positive markers and, optionally, genes expected to be absent. Genes missing from the count matrix stop the run with an error before scoring. Scores are computed with a BPCells-based implementation of UCell scoring (Andreatta and Carmona 2021, Parks and Greenleaf 2025). It processes the raw GEX counts in blocks of limited size and reproduces UCell's per-cell descending ranks, rank truncation at 1,500 genes, positive and negative signatures and lower-bound clipping. Working in blocks avoids loading the whole matrix or building a Seurat object only for scoring. The pipeline does not use the UCell package itself. Validation instead requires agreement with the UCell reference to within $10^{-12}$. Signed signatures are clipped at zero per cell before averaging. The same scoring annotates the GEX, ATAC and WNN clusters. WNN labels can use their own marker sets, whose matched controls come from the same GEX reference nuclei.

Raw UCell scores are not comparable between signatures, because some gene sets score highly by chance. Each label's cluster-level mean score is therefore compared with 999 random control signatures matched on gene abundance and detection. The control reference samples up to 50 nuclei per GEM well from the GEX metadata before doublet filtering. The GEX, ATAC and WNN annotations thus share the same controls, whatever nuclei each stage keeps. For each control signature, the markers are taken in random order. Each is replaced by a gene drawn from its 50 nearest eligible candidates not yet used in that signature. Candidates exclude all marker genes and undetected genes. A label's adjusted score in a cluster is its observed mean minus the 95th percentile of its matched controls. The label with the highest adjusted score is the candidate. Its advantage is the smaller of its lead over zero and its lead over the runner-up. The candidate is assigned when its advantage is positive and at least the configured minimum (0.05 by default). Otherwise, the cluster remains unassigned, with the candidate and reason kept. Requiring a lead over both the background and the runner-up leaves ambiguous clusters unassigned rather than forcing a label. Raising the minimum can therefore only make assigned clusters unassigned, never relabel them. Each unassigned cluster keeps its own label, `Unassigned_cluster_<id>`, so doublet detection, pseudobulks and per-cell-type tests never pool unrelated unassigned clusters. Because annotation depends on the supplied signatures and their level of detail, labels can represent either cell types or broader groups of cells, such as a lineage or a tissue of origin.

Annotation outputs also report, for each cluster, how many positive markers are detected in at least 10% of its nuclei.

#### Doublet removal

After GEX clustering, `scDblFinder` is applied per GEM well, with `dbr.sd = 1`, to the gene-expression counts of the kept nuclei (Germain *et al.* 2022). These counts are CellBender-corrected where configured. Doublets form only within a GEM well, and `dbr.sd = 1` sets the doublet threshold from misclassification alone rather than from an expected doublet rate. scDblFinder receives cluster labels from the annotation: the cell-type label of an assigned cluster, or otherwise its annotation status and identifier. A configured map can optionally collapse these labels. By default, nuclei called as doublets are removed, as are all nuclei of Leiden clusters whose doublet fraction exceeds 0.5.

### ATAC processing

#### Peaks and LSI

Fragments on the standard chromosomes are used for peak calling. Peak-calling groups are defined by a configurable metadata column of the nuclei kept after GEX processing, by default their GEX clusters. GEX clusters serve because ATAC clustering needs the peak matrix. Calling peaks per group keeps those of rare populations, which a pooled call would dilute (Granja *et al.* 2021, Stuart *et al.* 2021). Groups larger than 50,000 nuclei are randomly downsampled for peak discovery, which limits the runtime and memory of peak calling. Each group's discovery nuclei are split into two random halves. Peaks are called in each half with either MACS3 or the BPCells tile-based caller (Zhang *et al.* 2008, Parks and Greenleaf 2025). MACS3 runs on exported group fragments with a −75-bp shift, a 150-bp extension and summit calling. The tile-based caller runs on the stored fragments, using 500-bp tiles and an FDR of 0.01. MACS3 reads each exported fragment as a single read at its left end, so only one of its two insertions enters the pileup. Consensus peaks follow the ArchR strategy of iterative fixed-width peak construction (Granja *et al.* 2021). Summits are extended by 250 bp to 500-bp peaks, and peaks overlapping any blacklisted region of the reference genome are removed (Amemiya *et al.* 2019, Ogata *et al.* 2023). Overlapping peaks are collapsed by summit significance, first within each half and then across the two halves. Only peaks called in both halves are kept. Like ArchR's pseudo-replicate rule (Granja *et al.* 2021), this removes weak peaks called in only one half. Each group keeps its strongest peaks, up to 500 per discovery nucleus and 150,000 in all, as ArchR does by default (Granja *et al.* 2021). The group-level sets are then combined into one non-overlapping consensus set. Here, overlapping peaks are ranked by the quantile of their summit significance within their group. The caps and within-group quantiles keep large or deeply sequenced groups from dominating the consensus (Corces *et al.* 2018). BPCells counts fragment insertions (by default) or overlaps within these peaks for all nuclei kept after GEX processing. Insertions, which ArchR also counts, measure Tn5 cuts inside each peak, whereas overlaps also count fragments that merely span it.

ATAC dimensional reduction starts from the consensus peak matrix of the nuclei that pass the aggregation-level ATAC exclusions. Signac's TF-IDF method 1, with a scale factor of 10,000, normalizes this matrix (Stuart *et al.* 2021). A BPCells singular-value decomposition of the result gives the latent semantic indexing (LSI) embeddings (Parks and Greenleaf 2025). The pipeline keeps all LSI components and does not test them for correlation with sequencing depth. Instead, the default dimension settings leave out the first component, which often reflects depth (Stuart *et al.* 2021).

#### Filtering and doublet removal

ATAC processing starts from the nuclei kept after GEX processing. The peak-calling groups and the consensus peak matrix are built on this cell set. Per-nucleus quality-control metrics, such as the counts in peaks and their fraction, are calculated from the matrix and checked against configured aggregation-level exclusions. After ATAC clustering, a separate ATAC-based `scDblFinder` analysis follows per GEM well, with `dbr.sd = 1`. Instead of scDblFinder's internal per-well feature aggregation, peaks are grouped once into 50 feature groups. The groups come from *k*-means clustering of the peaks' aggregation-wide LSI coordinates on dimensions 2–20 (loadings scaled by the singular values, as in scDblFinder). The peaks of each group are summed with BPCells before scDblFinder's unchanged classifier runs. This change limits memory use, but its scores are not expected to equal those from the internal aggregation. That aggregation transforms every peak within each well, which can exhaust memory before classification. ATAC doublets are removed at the nucleus and cluster level as for GEX.

##### Motif accessibility

Transcription-factor motifs are represented by the 233 sequence-similarity families of the JASPAR 2026 CORE vertebrate collection (Ovek Baydar *et al.* 2026). Families are used because related transcription factors share near-identical motifs, whose matches and accessibility cannot be told apart. The pipeline includes a copy of the familial root motifs and the family membership table. Each root motif is trimmed of flank columns whose total count is less than half of its largest column total. These columns come from a minority of the family's aligned member motifs, so their counts are sparse and can look specific by chance. Scanning with them would restrict matches to that minority's sub-family. The trimmed root motifs are matched to the consensus peaks with `motifmatchr` at its default threshold of $P < 5 \times 10^{-5}$ (Schep A 2025). Configured transcription factors are matched to families by name, or by name and motif identifier when a symbol belongs to more than one family. betterChromVAR applies the chromVAR workflow with GC-bias correction to compute analytic motif-family deviations and z-scores for individual nuclei (Schep AN *et al.* 2017, Germain 2026). All families are tested per ATAC annotation class with one-versus-rest Wilcoxon tests, with Benjamini–Hochberg correction (Benjamini and Hochberg 1995) across all family–class tests. They are also summarized as cell-weighted means per ATAC cluster, GEX cluster and GEX cell type. The configured transcription factors select only the families shown on embeddings. These values measure accessibility associated with a motif family and are not direct measurements of transcription-factor activity.

#### Weighted nearest-neighbor integration

GEX and ATAC representations are combined with a BPCells-based implementation of the Seurat weighted-nearest-neighbor (WNN) strategy (Hao *et al.* 2021, Parks and Greenleaf 2025). The native implementation returns neighbor indices, distances and modality weights that clustering, UMAP and SCAVENGE reuse, without building a Seurat object. The GEX and ATAC embeddings, after optional Harmony correction, are aligned by barcode and L2-normalized. For each modality separately, 200 candidate neighbors are then found with hierarchical navigable small world (HNSW) search (Malkov and Yashunin 2020). multiomeR sets each cell's kernel width in each modality from the configured neighbor count, following Seurat's small-SNN bandwidth strategy. Per-cell modality weights are computed from these kernels by comparing within-modality with cross-modality predictions. The candidates of both modalities are then pooled, and those with the highest weighted kernel scores become the final neighbors. In our tests, Seurat's weights approached 0 or 1 as the number of nuclei grew, because the kernel widths shrink with density. The weights are therefore estimated within random subsets of at most 10,000 nuclei, whereas the neighbors are selected among all nuclei. BPCells builds an SNN graph from the joint neighbors for Leiden clustering, and UMAP uses the joint neighbor lists directly. The implementation does not use `Seurat::FindMultiModalNeighbors()`. Because HNSW replaces Seurat's Annoy search, exact equality is not expected. Its validation instead requires the modality weights to correlate with Seurat's, and the neighbors to overlap with Seurat's, above preset thresholds.

WNN analysis uses the nuclei kept after ATAC processing that are present in both embeddings. When small joint clusters are removed, the neighbors and modality weights are recomputed on the kept nuclei, while the cluster labels are kept. The UMAP is computed once, from the final neighbors. The GEX, ATAC and WNN metadata therefore describe successively smaller sets of nuclei, not interchangeable versions of the Cell Ranger cell calls.

### Differential analyses

#### Cell-type composition

The differential-analysis module treats donors, rather than nuclei, as the biological replicates. It takes annotation classes from the WNN cell-type label of the final WNN metadata. For cell-type composition, nuclei are counted by donor and annotation class, adding missing combinations as zero counts. Unless the model lists the response classes, the assigned classes observed in at least half of the donors are tested. Rarer classes and unassigned clusters are not tested, as they would only add to the multiple-testing burden, but they still count in every donor's total. Each class is modeled separately in a fixed-effects beta-binomial model with logit link, fitted by `glmmTMB` with the configured formula (Brooks *et al.* 2017). Its response has two columns: the numbers of nuclei in that class and in all other classes. Named linear contrasts of the fixed effects are tested with Wald statistics. Fits that fail, do not converge or lack a positive-definite Hessian are reported as non-estimable. Benjamini–Hochberg correction is applied across the tested annotation classes within each model and contrast, but not across contrasts or models.

#### Molecular pseudobulk analyses

For molecular analyses, GEX and ATAC counts of the final WNN nuclei are summed within each donor–annotation-class combination to form pseudobulk samples. Four feature matrices are tested:

- **Gene expression** (differential gene expression, DGE): gene pseudobulk counts.
- **Chromatin accessibility** (differential chromatin accessibility, DCA): consensus-peak pseudobulk counts.
- **Motif-family accessibility** (differential transcription-factor motif accessibility, DTFA): betterChromVAR bias-corrected deviations of the 233 JASPAR motif families, calculated from the pseudobulk ATAC counts (Germain 2026). This matrix is therefore continuous. Unlike chromVAR $z$-scores, deviations do not grow with sample depth, and no per-sample centering or cross-sample normalization is applied.
- **Transcription-factor activity** (differential CollecTRI transcription-factor activity, DCTA): signed CollecTRI regulator activities, inferred with the `decoupleR` univariate linear model from filtered, library-size-normalized log-CPM GEX pseudobulks (Badia-i-Mompel *et al.* 2022, Müller-Dott *et al.* 2023). Each regulator requires at least five measured targets. The CollecTRI network is used only when its checksum matches the recorded one, and complexes such as AP1 and NF-κB are kept as single regulons.

For each model, donors that miss a model variable or fall outside an optional donor list are removed. Samples outside an optional annotation-class subset are also removed, and the DTFA analysis by default removes samples with fewer than 5,000 ATAC counts. For count matrices, zero-depth samples are removed, features are filtered with the design-aware `edgeR::filterByExpr()` (Chen *et al.* 2025), and library sizes are TMM-normalized with `edgeR::normLibSizes()` (Robinson *et al.* 2010, Robinson and Oshlack 2010).

Model matrices are built from a configured formula or custom design function, and contrasts from named linear expressions or custom contrast functions. A configured random effect selects `limma::duplicateCorrelation()` with the random effect as block (Smyth *et al.* 2005, Ritchie *et al.* 2015). For counts, voom precision weights (Law *et al.* 2014) are re-estimated for the block and correlation (`edgeR::voomLmFit()`). Wherever an intra-block correlation is estimated, the consensus is taken from at most 50,000 evenly spaced features, enough to estimate this one value pooled across all features (Smyth *et al.* 2005) at a fraction of the runtime. The models are still fitted to all features. Without a random effect, count designs with more than one coefficient are fitted with edgeR quasi-likelihood models and tested with quasi-likelihood F-tests (Lund *et al.* 2012, Chen *et al.* 2025). These models use robust empirical Bayes moderation of the dispersions (Phipson *et al.* 2016). Continuous matrices and single-coefficient designs use limma. A single-coefficient count design is fitted on the raw counts, without the filtering and normalization above. An optional paired-cell-type route fits each annotation class separately with voom–limma, or limma for continuous matrices. It allows at most one pseudobulk per pairing unit and class. A cross-class contrast is tested on a combined response, restricted to the donors present in every class involved. This response is the contrast-weighted sum of the classes' log-expression. It is weighted by the inverse of its variance, which is computed from the classes' precision weights. The combined response is fitted with the same donor-level design and correlation block, so each feature has its own covariance between classes.

For each model and contrast, Benjamini–Hochberg FDR is calculated across the tested features, and features with an FDR below 0.05 are significant. DGE contrasts are also tested against the MSigDB Hallmark and Reactome collections (Liberzon *et al.* 2015, Milacic *et al.* 2024) with competitive `camera` tests (Wu and Smyth 2012, Dolgalev 2026). These tests use the contrast's expression, precision weights and design, or the combined response for cross-class contrasts. Each set's inter-gene correlation is estimated from the residuals, and each set needs at least ten tested genes. FDR is calculated within each contrast and collection. A cross-modality comparison maps CollecTRI regulators to JASPAR families. It compares the model $t$-statistics, rather than the raw scales, of transcription-factor activity and motif-family accessibility. Where gene expression is fitted with limma, it also compares TF expression (edgeR tests report no $t$-statistic). Family-level summaries use the median regulator $t$-statistic. An analysis stops with an error rather than returning a model when no samples or testable features remain or residual degrees of freedom are below one. It also fails when a correlation block or contrast is invalid, or when too few shared donors remain for a requested paired-cell-type comparison.

### Peak–gene correlation

The optional peak–gene correlation module runs separately in each assigned WNN annotation class of at least 200 nuclei. It requires donor identifiers and per-nucleus GEX and ATAC depth metadata. Classes without them are skipped, and the reason is recorded. Consensus ATAC peaks are paired with gene transcription start sites (TSSs) on the same chromosome within 250 kb of the peak center. Each pair is classified as self-promoter (−1,500 to +500 bp around the TSS), gene-body, proximal (within 10 kb) or distal. Within each class, nuclei of donors contributing at least 40 nuclei are split into mutually exclusive ATAC-state bins. The bins come from $k$-means clustering of scaled ATAC LSI dimensions 2–20, Harmony-corrected where configured. The number of bins is the median number of nuclei per donor divided by 20, rounded down and capped at 20. Counts are summed within each donor–state combination to create non-overlapping pseudobulks. Unlike ArchR's overlapping nearest-neighbor aggregates, which reuse nuclei, disjoint donor–state pseudobulks keep samples independent and donors identifiable (Granja *et al.* 2021). Pseudobulks of fewer than 20 nuclei are dropped. Donors and states are then removed in repeated rounds until every donor contributes at least two states.

GEX and ATAC pseudobulk counts are separately scaled to counts per million, using the pseudobulk depth from the metadata, and log1p-transformed. Genes and peaks must be detected in at least 5% of pseudobulks. Each chromosome's analysis requires at least 10 pseudobulks and 5 residual degrees of freedom. A measurement-support filter then removes peak–gene pairs whose gene and peak are not both supported. Like `edgeR::filterByExpr()`, it drops pairs too sparsely measured for reliable inference, which also cuts runtime and the multiple-testing burden (Chen *et al.* 2025). Its count thresholds are scaled by each pseudobulk's depth relative to the class median, but never fall below 2. The gene and the peak must each reach their thresholds in at least the larger of a minimum number and a fraction of the pseudobulks. They must also do so in at least two pseudobulks of each of a minimum number of shared donors. The lenient preset requires RNA and ATAC counts of at least 5 and 3, the larger of 6 pseudobulks and 10% of them, and two shared donors. The default moderate preset raises the counts to 10 and 5. The strict preset also requires the larger of 10 pseudobulks and 20% of them, and three shared donors with three supporting pseudobulks each. Excluded pairs are never tested and do not enter the multiple-testing family.

GEX and ATAC values are residualized against donor, scaled log RNA and ATAC depth and the class's accessibility components described below. The Pearson correlation of the residuals is reported as a descriptive summary. The hierarchical analysis fits donor fixed intercepts, depth covariates, the accessibility components and a donor-varying slope for the within-donor-centered peak value. It uses profiled restricted maximum likelihood, with Kenward–Roger inference for the average slope (Kenward and Roger 1997). The donor-varying slope makes donors rather than pseudobulks the units of inference. Kenward–Roger adjusts its standard error and degrees of freedom for the small number of donors. A peak is tested only when its accessibility varies within at least eight donors. With fewer, the random-slope test has too few degrees of freedom to be informative, so its pairs remain untested. Fits that fail numerical checks keep their estimates but get no P-value. Peaks and genes that both vary along shared chromatin states correlate whether or not they interact. The accessibility components remove the leading states. They are the top ten principal components, or one per ten residual degrees of freedom when fewer. They are computed from all candidate peaks, across chromosomes, that are accessible in at least 5% of the class's pseudobulks. Each peak enters as its donor- and depth-residualized, standardized log1p CPM. Genes that vary along the remaining states still correlate with many peaks. Each candidate is therefore compared with its own gene's background, like Signac's comparison of each peak–gene correlation with matched background peaks (Stuart *et al.* 2021). This background consists of up to 200 candidate peaks that lie on the same chromosome but more than 10 Mb from the gene's TSS, scored with the same model. The candidate's signed normal score is the normal quantile of its P-value, signed by the slope. This score is standardized by the mean and SD of the gene's background scores, given at least ten of them. It is then rescaled by the class's background SD within its peak-detection quintile. A candidate's empirical P-value is the share of the class's rescaled background scores that are at least as extreme. These P-values are Benjamini–Hochberg-corrected within each annotation class over the complete eligible pair family, counting untested and unreliable pairs. A candidate link requires an estimable positive average slope above the gene's background mean and an FDR below 0.05, excluding self-promoter peaks. SuSiE fine-mapping prioritizes peaks on the same residualized values for genes with at least one candidate link (Wang *et al.* 2020). Because neighboring peaks are often co-accessible (Pliner *et al.* 2018), SuSiE asks which of a gene's correlated peaks jointly explain its expression. It fits up to ten single effects and reports 95% credible sets whose peaks have a minimum absolute correlation (purity) of at least 0.5. Fine-mapping covers up to the 50 genes with the lowest FDR per class and chromosome, and the 500 peaks per gene with the largest absolute residual correlation. These caps limit its runtime. These model-based associations do not establish causal enhancer–gene regulation. Background peaks more than 10 Mb away also cannot separate a specific enhancer from covariation within a local chromatin domain.

Top-link plots rank estimable positive hierarchical associations by their uncalibrated Kenward–Roger P-value, without a significance cutoff. They exclude self-promoter peaks but keep gene-body peaks. Each plot shows the gene context, the plotted class's insertion coverage and the donor-residual scatter.

### Genetic enrichment

Each multiomeR version uses one fixed Open Targets Platform release, currently 26.03, and downloads its study, credible-set, credible-set evidence and target datasets (Buniello *et al.* 2025). Users can supply their own fine-mapping results as Parquet files in a fixed format. For each configured GWAS, multiomeR retrieves the requested study and fine-mapping result. When selection is automatic, it takes the first available of SuSiE, SuSiE-inf and PICS fine-mapping (Farh *et al.* 2015, Wang *et al.* 2020, Cui *et al.* 2024). Variants of the study's credible sets (95% sets for Open Targets) with a posterior probability above 0.001 by default are mapped to consensus ATAC peaks. Variant weights that map to the same peak are summed and capped at one. These peak weights form the trait annotation from which analytic nucleus-level chromVAR z-scores are calculated, reusing the GC-bias background of the motif analysis (Schep AN *et al.* 2017, Ulirsch *et al.* 2019, Germain 2026). The background variance of a weighted sum of peaks scales with the squared weights. betterChromVAR instead scales the variance of its analytic z-scores with the summed weights, so the z-scores are rescaled accordingly. To show which traits are not independent, credible sets are also compared pairwise. Posterior probabilities are summed per variant and normalized to one within each GWAS. The similarity of two GWAS is the sum, over identical variants, of the smaller of their two values. It equals one minus the total variation distance and is not a colocalization test.

Annotation-class pseudobulk deviations are calculated separately rather than by averaging the nucleus-level results. ATAC counts are summed by the final WNN cell-type labels, which the other analyses also use. A betterChromVAR background model is fitted to the resulting peak-by-class matrix. The expected accessibility of each peak averages the depth-normalized class profiles with equal weight. Each class is thus compared with the average class rather than with a reference dominated by the most abundant classes. Background peaks are matched on this expectation and on GC and fragment-length bias. The raw deviation is the observed-minus-background accessibility of the weighted peaks relative to their expected accessibility. A relative deviation also standardizes the raw deviations across annotation classes within each trait. Analytic z-scores, rescaled for the peak weights as for nuclei, give one-sided P-values. Their Benjamini–Hochberg-adjusted values across all traits and annotation classes are reported. Plot labels mark unadjusted z-scores of at least 1.645 and 2.326. When effect sizes are available, an absolute-effect version weights variants by posterior probability times absolute effect size. When only loci carry effects, each locus's lead-variant effect is used. These weights are rescaled to sum to the kept variants' posterior probabilities. Each class deviation is split into additive contributions from its loci and variants, which sum to the class totals. The loci and variants are labeled with the highest-scoring Open Targets locus-to-gene genes (Mountjoy *et al.* 2021). Detail plots are drawn for classes with a positive deviation and a z-score of at least 1 by default, in either weighting. They show the top loci by absolute contribution and by combined contribution and effect-size rank.

SCAVENGE-style trait-relevance scores are calculated from the nucleus-level z-scores on the WNN graph. They use multiomeR's own sparse-matrix implementation of the SCAVENGE propagation strategy rather than the reference package (Yu *et al.* 2022). The reference package's dependencies predate the pipeline's R environment, and the local version propagates directly over the sparse WNN graph. Nuclei whose one-sided normal-tail probability is at most 0.05 are seeds. When they exceed the maximum seed fraction, 5% of nuclei by default, those with the highest z-scores are kept. When no nucleus qualifies, all scores are zero. The WNN graph is made unweighted, with every nonzero edge set to one, and nuclei without edges are excluded. Seed signal is propagated by a random walk with restart probability 0.05 by default until convergence. Nuclei that no seed reaches remain unscored. Scores are capped at their 95th percentile, min–max scaled and multiplied by the mean z-score of the top 1% of nuclei. Unlike the reference, the pipeline computes no permutation P-values. The reference's degree-matched null scatters seeds across the graph, whereas observed seeds cluster within cell types. Nearly every seeded group would therefore appear significant. Scores are summarized per metadata group.

#### Settings for the example application

The example application used the `mixed_human_31x` aggregation of `configuration_dev/` in the archived release. The settings below include the defaults it left unchanged.

##### Inputs and quality control

The aggregation combined the 31 samples of Supplementary Table S2, reprocessed from the raw reads with Cell Ranger ARC 2.1.0 (10x Genomics 2026) and the GRCh38 2024-A reference. Their GEX counts were corrected for ambient RNA with CellBender 0.4.0 `remove-background` (Fleming *et al.* 2023). CellBender ran with default settings on the raw Cell Ranger matrix without the peak features. Every sample excluded nuclei with `amulet_q.value < 0.01`, `nCount_RNA < 250`, `nucleosome_signal > 4` or `TSS.enrichment < 4`, and those with `RNA_mito_percent > 10` (25 samples) or `RNA_mito_percent > 20` (6 samples). After peak calling, the aggregation also excluded nuclei with `nCount_ATAC < 1000`, `atac_peak_counts_frac < 0.1` or `atac_peak_counts_blacklist_frac > 0.05`.

##### Processing

Gene expression was normalized with the BPCells-native backend without regressing out covariates. The neighbor graphs and UMAP used GEX principal components 1–30 and ATAC LSI dimensions 2–30. Harmony batch correction was not used. The GEX, ATAC and WNN graphs used 50 neighbors and Leiden resolutions of 0.8, 0.8 and 0.4, respectively, with a minimum cluster size of 50 nuclei. UMAP used 50 neighbors and `min_dist = 0.001`. Nuclei that scDblFinder called as GEX or ATAC doublets were removed, as were clusters with a doublet fraction above 0.5. Peaks were called with MACS3 in each GEX cluster, and the consensus peak matrix counted fragment insertions.

##### Cell-type annotation

The GEX signatures were B (`MS4A1`, `CD79A`); T (`TRAC`, `CD3D`); Mono (`LYZ`, `S100A9`); NK (`KLRD1`, `GNLY`); pDC (`LILRA4`, `CLEC4C`); Plasma (`TNFRSF17`, `SDC1`, `DERL3`); Inhibitory neuron (`GAD1`, `GAD2`, `SLC6A1`, `SNAP25`); Oligodendrocyte (`MBP`, `PLP1`, `MOG`, `MAG`); Microglia (`P2RY12`, `TMEM119`, `SALL1`); Macrophage (`CD163`, `MRC1`, `FOLR2`, `MSR1`); Pancreatic beta (`INS`, `IAPP`, `PCSK1`); Pancreatic acinar (`PRSS1`, `REG1A`, `CELA3A`); Ductal epithelial (`CFTR`, `SCTR`, `PKHD1`, `KRT7`); Hepatocyte (`ALB`, `APOA1`, `CYP3A4`); Cardiomyocyte (`TNNT2`, `MYH6`, `MYH7`, `ACTC1`, `MYBPC3`, `TNNI3`, `TTN`, `RYR2`, `PLN`); Colonocyte (`CA1`, `KRT20`, `SLC26A3`); Goblet (`MUC2`, `SPINK4`, `CLCA1`); Alveolar type 1 (`AGER`, `CAV1`, `EMP2`); Alveolar type 2 (`SFTPC`, `SFTPA1`, `ABCA3`); Airway secretory (`SCGB1A1`, `SCGB3A1`, `SCGB3A2`, `CYP2F1`); Endothelial (`PECAM1`, `VWF`, `EMCN`); Fibroblast (`COL1A1`, `DCN`, `COL3A1`); Smooth muscle (`ACTA2`, `TAGLN`, `MYH11`); Pericyte (`RGS5`, `PDGFRB`, `CSPG4`). For the WNN labels, an islet endocrine signature (`INS`, `SLC30A8`, `PCSK2`, `ABCC8`, `KCNMB2`) replaced the Pancreatic beta signature, which tied with the acinar signature on ambient `INS` in the islet cluster. The minimum annotation advantage was 0.05.

##### Peak–gene correlation and genetic enrichment

The peak–gene correlation module used its default settings, including the moderate measurement-support filter. The genetic enrichment module tested the 26 Open Targets studies of Supplementary Table S3. They come from GWAS of traits related to the sampled tissues and blood. The number of effective peaks of a study is $(\Sigma w)^2/\Sigma w^2$ of its peak weights $w$. Only studies with at least 20 effective peaks in this aggregation were kept, so that no study rests on one or a few loci. A fasting-glucose study (GWAS Catalog GCST90278658, Cerezo *et al.* 2025) was also left out because another study in Supplementary Table S3 (Jee *et al.* 2025) contains its samples. Fine-mapping methods were selected automatically, with a posterior-probability cutoff of 0.001. SCAVENGE used a restart probability of 0.05 and a maximum seed fraction of 0.05.

#### Algorithmic reimplementations and validation

Four published algorithms were reimplemented so that matrices stay on disk or sparse and steps run in parallel. They are UCell scoring, AMULET, WNN and SCAVENGE propagation, each described with its analysis step above. The ATAC feature aggregation before scDblFinder is an adaptation rather than a reimplementation. Automated tests compare each with its reference software or an independent calculation. They require exact equality where expected and predefined similarity otherwise. The *Algorithm validation* page of the user manual lists the intended differences and how to rerun each check.

Supplementary Table S4 compares each implementation with its reference on objects of `comparison_5x`, the first five samples of Supplementary Table S2. For UCell, it measures a nucleus-level scoring function rather than the cluster-level function that annotation uses. The automated tests compare that function with UCell to within $10^{-12}$. For WNN, it estimates the modality weights from all nuclei, whereas the pipeline estimates them in random subsets of at most 10,000 nuclei. It therefore covers the weighting and neighbor selection but not the subsetting. Each implementation ran once, as a separate pipeline step, in a new process on a six-core Slurm job. A new process keeps memory left over from earlier steps out of the peak RAM. Wall time includes data loading, package setup, data conversion and compilation. Peak RAM is the maximum sampled resident set size of the process tree. The AMULET reference used scDblFinder's full-memory mode, which runs serially, so that both implementations read the same complete input. scDblFinder's default per-chromosome mode was not compared. The ATAC scDblFinder input matches the pipeline's data before doublet removal, with insertion-counted consensus peaks and no cluster labels. With the cluster labels that the pipeline supplies, the score-rank Spearman correlation was 0.677, the doublet-call Jaccard index 0.817 and the class agreement 97.93%. The SCAVENGE reference ran the SCAVENGE 1.0.2 propagation functions on the same graph and seed z-scores. The comparison therefore covers propagation but not how the graph is built. Because `comparison_5x` runs no genetic enrichment, the chromVAR z-scores of a motif family were used as seeds. These single-run comparisons are descriptive, not a formal performance study.

#### Benchmark against a Seurat/Signac workflow

Supplementary Figure S2 compares multiomeR with a Seurat v5.6/Signac v2 baseline (Stuart *et al.* 2021, Hao *et al.* 2024) on nested aggregations of the samples of Supplementary Table S2. The aggregations are the first 1, 2, 5, 10 and 20 samples and all 31 samples (2,713 to 375,757 nuclei). Both workflows ran from the Cell Ranger ARC outputs to joint WNN clustering and embedding. Neither used per-sample quality-control filters, CellBender or doublet removal, so both kept the same nuclei and called the same consensus peaks. The code of both workflows and of the measurements is in `pipeline_manuscript_figures/` of the archived release.

The baseline is a short targets pipeline that runs a standard Seurat v5.6/Signac v2 analysis, written by the authors. It uses Seurat 5.6.0, the Signac v2 development version that multiomeR uses (1.9999.12) and fragtk 1.7.0. It used the same Slurm job types as multiomeR, with the same dimensions, neighbor counts and clustering resolutions. It keeps gene-expression counts and Signac's FeatureMatrix peak counts in BPCells directories, and calls peaks with MACS3 on multiomeR's GEX clusters with the same consensus-peak steps. Otherwise it runs the standard SCTransform, PCA, TF-IDF, LSI, WNN, UMAP and clustering steps. Seurat's multithreaded functions used the six cores of each Slurm job, and peak calling and counting ran one process per core through the future package (Bengtsson 2021).

Like the baseline, multiomeR normalized gene expression with Seurat SCTransform rather than its default BPCells backend, so that both workflows normalized identically. It was measured up to its multimodal Seurat/Signac export, the counterpart of the baseline's final Seurat object. Its measurements exclude analyses that the baseline does not perform. These include cell-type annotation, doublet detection, motif analysis and the extra embeddings. The shared preparation of cluster-level fragment inputs was excluded from both workflows.

The critical path of multiomeR is the longest chain of dependent steps, measured as the sum of their recorded runtimes, assuming that all independent steps run at the same time. The critical path of the baseline, which a script would run step by step, is the sum of its steps' runtimes. Neither is observed wall time, which on a shared cluster also includes queueing and job start-up. CPU time and RAM use sum each step's runtime multiplied by its mean CPU and memory use. CPU and memory use were sampled every 10–15 s from each step's Slurm control group, excluding reclaimable page cache. Steps too short to sample were assigned the median of the sampled steps. Peak RAM is the largest sampled memory use of a single step. Disk space, in decimal GB, sums the saved pipeline outputs and the Cell Ranger outputs that both workflows read. These are the filtered matrices, fragment files and their indexes (0.5 to 67.2 GB).

Each workflow ran once, multiomeR in an earlier run of the same pipeline with Seurat 5.5.1. On `comparison_5x`, multiomeR's SCTransform step gave identical principal components and took 289–322 s with either Seurat version. Supplementary Table S6 lists every measure at each size. At one sample, multiomeR used similar CPU time but more memory gigabyte-hours (0.30 versus 0.16 GB-hours). It used more disk space at every size, mainly because it keeps each sample's fragments in BPCells format, a copy that every aggregation of that sample reuses.
