## Supplementary Tables S1-S6 for "multiomeR: a pipeline for reproducible analysis of large-scale single-nucleus Epi Multiome datasets"

### multiomeR: a pipeline for reproducible analysis of large-scale single-nucleus Epi Multiome datasets

Thomas Gade Koefoed[1, *], Shanshan He[1], Malte Thodberg[1], Laura Maarit Pikkupeura[1], Kata Krizic[1], Benedicte Schultz Kapel[1], Raquel Sanz Martinez[1], Simon Rasmussen[1], Torben Hansen[1, *]

[1] Novo Nordisk Foundation Center for Basic Metabolic Research, Faculty of Health and Medical Sciences, University of Copenhagen, Blegdamsvej 3B, 2200 Copenhagen N, Denmark

**This file includes:**

Supplementary Table S1: Workflow characteristics and analyses of selected approaches to Epi Multiome studies after Cell Ranger ARC
Supplementary Table S2: Public 10x Genomics and ENCODE Epi Multiome datasets used for the example application (`mixed_human_31x`)
Supplementary Table S3: GWAS used in the example application of the genetic enrichment module
Supplementary Table S4: Agreement and resource use of multiomeR's reimplementations and their reference implementations
Supplementary Table S5: Main tools, packages and reference resources used by multiomeR
Supplementary Table S6: Resource use of multiomeR and a Seurat/Signac baseline at each aggregation size

#### Supplementary Table S1

**Workflow characteristics and analyses of selected approaches to Epi Multiome studies after Cell Ranger ARC.** Categories summarize documented support in the linked software versions: Native means provided by the approach, Limited means partial support (see notes), and External/custom means left to other tools or user code.

| | **multiomeR**<br>1.2.2 | **Seurat + Signac**<br>5.6.0 + 1.9999.12 | **ArchR**<br>1.0.3 | **SnapATAC2 + muon**<br>2.10.0 + 0.2.0 | **SCENIC+**<br>1.0a2 |
|---|---|---|---|---|---|
| **Workflow** | | | | | |
| Project-level multi-sample configuration | **Native** | External/custom | Limited[a] | External/custom | External/custom[b] |
| Reruns only changed steps and records how results were made | **Native** | External/custom | External/custom | External/custom | **Native**[c] |
| Disk-backed core analysis | **Native** | Limited[d] | Limited[e] | Limited[f] | External/custom |
| Scheduler-aware parallel execution | **Native** | External/custom | External/custom | External/custom | **Native**[c] |
| **Analyses** | | | | | |
| Doublet detection in both modalities | **Native** | External/custom | Limited[g] | **Native**[h] | External/custom |
| Donor-level pseudobulk differential testing | **Native** | Limited[i] | External/custom | **Native**[j] | External/custom |
| Peak–gene links with donors as replicates | **Native** | Limited[k] | Limited[k] | Limited[k] | Limited[k] |
| Cell-type enrichment of fine-mapped GWAS variants | **Native** | External/custom | Limited[l] | External/custom | External/custom |
| Nucleus-level GWAS enrichment | **Native** | External/custom | External/custom | External/custom | External/custom |

**Versions and sources.**
*multiomeR:* 1.2.2, commit `ca5d8f2`, built on `targets` (Landau 2021) and BPCells (Parks and Greenleaf 2025).
*Seurat + Signac:* Seurat 5.6.0 (Hao *et al.* 2024) and the Signac development version 1.9999.12 at commit e235116 (Stuart *et al.* 2021), run in R scripts.
*ArchR:* 1.0.3 (Granja *et al.* 2021), run as an ArchR project.
*SnapATAC2 + muon:* SnapATAC2 2.10.0 (Zhang K *et al.* 2024), muon 0.2.0 (Bredikhin *et al.* 2022) and other scverse packages (Virshup *et al.* 2023), run in Python scripts.
*SCENIC+:* 1.0a2 (Bravo González-Blas *et al.* 2023), run as a Snakemake workflow (Mölder *et al.* 2021).

**Notes.**
[a] An ArchRProject links sample Arrow files, but project setup, metadata and parameters remain defined in user-authored R code.
[b] SCENIC+ starts from a processed pycisTopic object (Bravo González-Blas *et al.* 2023) and a processed scanpy AnnData file (Wolf *et al.* 2018), so samples are combined in upstream scripts.
[c] Through Snakemake, which by default reruns jobs whose code, inputs, parameters or software environment changed, and submits jobs to schedulers such as Slurm through executor plugins.
[d] Seurat v5 and the Signac development version support BPCells-backed assay layers, including peak counts. However, Seurat 5.6.0's SCTransform (Hafemeister and Satija 2019, Choudhary and Satija 2022) loads the count layers into memory and returns dense residuals for all nuclei, so its memory use grows with the number of nuclei.
[e] HDF5 Arrow files back ATAC fragments, matrices and results computed from them. RNA and multimodal integration depend on the chosen route.
[f] SnapATAC2 provides fully backed AnnData and AnnDataSet objects for ATAC. RNA and multimodal backing depends on the selected scverse components and operations. muon uses HDF5-backed MuData and AnnData storage, but in-memory behavior depends on the selected analysis method.
[g] ArchR infers doublets from scATAC-seq data only.
[h] SnapATAC2 and scanpy provide Scrublet (Wolock *et al.* 2019) for ATAC and gene-expression counts, respectively.
[i] Seurat sums counts per donor and cell type with AggregateExpression and compares two groups with DESeq2 (Love *et al.* 2014) through FindMarkers, which takes no covariates for this test.
[j] pertpy tests decoupler pseudobulks with edgeR (Robinson *et al.* 2010) or PyDESeq2 (Muzellec *et al.* 2023) under design formulas that can include covariates (Badia-i-Mompel *et al.* 2022, Heumos *et al.* 2026).
[k] Signac's LinkPeaks and the network functions of SnapATAC2 (beta) correlate peaks with genes across nuclei, ArchR across groups of similar nuclei, and SCENIC+ ranks region–gene pairs by gradient-boosting importance and rank correlation. All provide peak–gene links, but none uses donors as replicates.
[l] ArchR computes chromVAR deviations (Schep AN *et al.* 2017) for user-supplied region sets, which can hold trait-associated variants.

#### Supplementary Table S2

**Public 10x Genomics and ENCODE Epi Multiome datasets used for the example application (`mixed_human_31x`).** ENCODE accessions resolve at https://www.encodeproject.org (ENCODE Project Consortium *et al.* 2020, Luo *et al.* 2020) and 10x Genomics dataset names at https://www.10xgenomics.com/datasets. Input nuclei are barcodes called as nuclei by Cell Ranger ARC 2.1.0 (10x Genomics 2026).

| Source identifier | Tissue and preparation | Input nuclei |
|---|---|---|
| `pbmc_granulocyte_sorted_3k` | PBMC; granulocytes removed | 2,713 |
| `10k_PBMC_Multiome_nextgem_Chromium_Controller` | PBMC; preparation not stated | 10,896 |
| `10k_PBMC_Multiome_nextgem_Chromium_X` | PBMC; preparation not stated | 11,152 |
| `pbmc_granulocyte_sorted_10k` | PBMC; granulocytes removed | 11,941 |
| `pbmc_unsorted_10k` | PBMC; unsorted | 12,220 |
| `pbmc_unsorted_3k` | PBMC; unsorted | 3,032 |
| `human_brain_3k` | Brain, cerebellum; 7-AAD-positive sorted nuclei | 3,454 |
| `ENCSR033MDU` | Pancreas | 5,165 |
| `ENCSR158DQA` | Pancreas | 18,567 |
| `ENCSR233SQG` | Pancreas | 12,051 |
| `ENCSR316WAS` | Pancreas | 6,340 |
| `ENCSR618WVK` | Liver | 17,570 |
| `ENCSR788DZO` | Liver | 8,487 |
| `ENCSR079UEO` | Heart | 34,779 |
| `ENCSR201BHG` | Heart | 16,438 |
| `ENCSR245BNT` | Heart | 16,002 |
| `ENCSR320RLS` | Heart | 12,793 |
| `ENCSR351CSQ` | Heart | 7,624 |
| `ENCSR508WRP` | Heart | 17,142 |
| `ENCSR511RDV` | Heart | 9,856 |
| `ENCSR534YJW` | Heart | 11,595 |
| `ENCSR568QKW` | Heart | 19,997 |
| `ENCSR613LBN` | Heart | 11,038 |
| `ENCSR793DNM` | Heart | 12,935 |
| `ENCSR923FKN` | Heart | 10,691 |
| `ENCSR956ZER` | Heart | 21,093 |
| `ENCSR987VTJ` | Heart | 12,427 |
| `ENCSR636WIB` | Colon | 6,396 |
| `ENCSR925IHI` | Colon | 14,464 |
| `ENCSR128ZLB` | Lung | 9,806 |
| `ENCSR264JIX` | Lung | 7,093 |
| **Total** | | **375,757** |

#### Supplementary Table S3

**GWAS used in the example application of the genetic enrichment module.** Fine-mapping methods and sample sizes were taken from Open Targets Platform release 26.03 (Buniello *et al.* 2025). Study IDs are GWAS Catalog accessions (Cerezo *et al.* 2025), each linked to its publication in PubMed. Sample size is the number of individuals in the discovery stage (cases plus controls). Supplementary Methods describes how the studies were selected. The final column reports, in order, the number of credible-set loci overlapping consensus peaks, unique credible-set variants overlapping those peaks, and unique consensus peaks receiving non-zero weight.

| Study ID | Trait | Method | Sample size | Credible-set loci | Overlapping loci / variants / peaks |
|---|---|---|---|---|---|
| GCST90002388 | Lymphocyte count (Vuckovic *et al.* 2020) | SuSiE-inf | 408,112 | 923 | 639 / 2,712 / 2,110 |
| GCST90002393 | Monocyte count (Vuckovic *et al.* 2020) | SuSiE-inf | 408,112 | 967 | 653 / 2,511 / 1,962 |
| GCST90446794 | Ulcerative colitis (Liu *et al.* 2023) | SuSiE-inf | 375,508 | 162 | 52 / 86 / 74 |
| GCST009597 | Multiple sclerosis (International Multiple Sclerosis Genetics Consortium 2019) | PICS | 41,505 | 140 | 113 / 573 / 442 |
| GCST90475159 | Body mass index (minimum, inverse-normal transformed) (Verma *et al.* 2024) | SuSiE-inf | 424,217 | 2,128 | 964 / 3,210 / 2,538 |
| GCST90105038 | Educational attainment (Okbay *et al.* 2022) | PICS | 3,037,499 | 1,598 | 922 / 3,747 / 2,886 |
| GCST90399686 | Asthma (Zhou *et al.* 2022) | SuSiE-inf | 1,376,071 | 159 | 105 / 340 / 288 |
| GCST90244092 | Lung function (FEV1) (Shrine *et al.* 2023) | PICS | 588,452 | 196 | 144 / 748 / 601 |
| GCST90244094 | Lung function (FEV1/FVC) (Shrine *et al.* 2023) | PICS | 588,452 | 318 | 231 / 1,134 / 874 |
| GCST90027158 | Alzheimer's disease (Bellenguez *et al.* 2022) | SuSiE-inf | 487,511 | 79 | 38 / 116 / 96 |
| GCST90128471 | Schizophrenia (Trubetskoy *et al.* 2022) | PICS | 175,799 | 222 | 154 / 792 / 646 |
| GCST90271609 | Genetic generalized epilepsy (International League Against Epilepsy Consortium on Complex Epilepsies 2023) | PICS | 59,945 | 20 | 13 / 58 / 47 |
| GCST003155 | Systemic lupus erythematosus (Bentham *et al.* 2015) | PICS | 14,267 | 41 | 34 / 204 / 146 |
| GCST90132222 | Rheumatoid arthritis (Ishigaki *et al.* 2022) | PICS | 276,020 | 96 | 75 / 405 / 303 |
| GCST90559230 | Atrial fibrillation (Roselli *et al.* 2025) | PICS | 1,650,345 | 307 | 226 / 1,226 / 906 |
| GCST007992 | Colorectal cancer (Law *et al.* 2019) | PICS | 92,967 | 66 | 51 / 195 / 154 |
| GCST90132314 | Coronary artery disease (Aragam *et al.* 2022) | SuSiE-inf | 1,165,690 | 245 | 158 / 582 / 470 |
| GCST90310294 | Systolic blood pressure (Keaton *et al.* 2024) | SuSiE-inf | 1,028,980 | 1,131 | 775 / 3,012 / 2,411 |
| GCST90310295 | Diastolic blood pressure (Keaton *et al.* 2024) | SuSiE-inf | 1,028,980 | 1,077 | 758 / 3,048 / 2,429 |
| GCST90301252 | Dupuytren's disease (Riesmeijer *et al.* 2024) | SuSiE-inf | 58,343 | 83 | 36 / 114 / 91 |
| GCST90239655 | LDL cholesterol (Graham *et al.* 2021) | PICS | 1,654,960 | 558 | 360 / 2,016 / 1,549 |
| GCST90239649 | HDL cholesterol (Graham *et al.* 2021) | PICS | 1,654,960 | 811 | 560 / 2,685 / 2,136 |
| GCST90239661 | Triglycerides (Graham *et al.* 2021) | PICS | 1,654,960 | 683 | 483 / 2,554 / 1,973 |
| GCST90428729 | Alanine aminotransferase (Ghouse *et al.* 2024) | PICS | 1,010,710 | 281 | 215 / 1,187 / 915 |
| GCST90492734 | Type 2 diabetes (Suzuki *et al.* 2024) | PICS | 2,535,601 | 687 | 507 / 2,336 / 1,850 |
| GCST90662896 | Fasting glucose (Jee *et al.* 2025) | PICS | 928,679 | 253 | 190 / 1,025 / 781 |

#### Supplementary Table S4

**Agreement and resource use of multiomeR's reimplementations and their reference implementations.** Each reimplementation and its reference ran once on the same input, taken from the first five samples of Supplementary Table S2 (all PBMC; 48,922 nuclei). The AMULET and ATAC scDblFinder inputs are single samples. The UCell row measures nucleus-level scoring, and the WNN row estimates modality weights from all nuclei (Supplementary Methods). Time and peak RAM are given for the reimplementation and then the reference, with their ratio in parentheses. A ratio below one means that the reimplementation used less. Supplementary Methods (Algorithmic reimplementations and validation) describes the references and measurements.

| Implementation | Input data | Reference | Result agreement | Time, reimplementation / reference | Peak RAM, reimplementation / reference |
|---|---|---|---|---|---|
| UCell scoring | 48,922 nuclei<br>38,606-feature GEX matrix | UCell 2.14.0 | Identical | 30.3 s / 76.4 s (0.40×) | 7.39 GB / 36.75 GB (0.20×) |
| AMULET | 12,220 nuclei<br>Complete fragment input | scDblFinder 1.24.10, full-memory AMULET | Identical | 25.5 s / 1,815.4 s (0.01×) | 0.87 GB / 29.69 GB (0.03×) |
| ATAC scDblFinder feature aggregation | 11,152 pre-doublet nuclei<br>159,533-peak matrix | scDblFinder 1.24.10 internal aggregation | Score-rank Spearman 0.777<br>Doublet-call Jaccard 0.811<br>Class agreement 96.82% | 31.9 s / 113.9 s (0.28×) | 1.31 GB / 10.59 GB (0.12×) |
| Weighted nearest neighbors | 48,922 nuclei<br>59-dimensional RNA/ATAC embeddings | Seurat 5.6.0 | RNA/ATAC weight Spearman 0.983 / 0.983<br>Mean neighbor overlap 0.965<br>First-quartile neighbor overlap 0.967 | 56.9 s / 62.5 s (0.91×) | 1.76 GB / 2.33 GB (0.75×) |
| SCAVENGE propagation | 48,922 nuclei<br>WNN graph | SCAVENGE 1.0.2 (`8ee8b173`) | Identical within floating-point precision | 6.7 s / 7.8 s (0.87×) | 0.67 GB / 0.69 GB (0.97×) |

**Sources.**
*UCell scoring:* UCell (Andreatta and Carmona 2021).
*AMULET and ATAC scDblFinder feature aggregation:* scDblFinder (Germain *et al.* 2022), whose full-memory mode runs AMULET (Thibodeau *et al.* 2021).
*Weighted nearest neighbors:* Seurat (Hao *et al.* 2021).
*SCAVENGE propagation:* SCAVENGE (Yu *et al.* 2022).

#### Supplementary Table S5

**Main tools, packages and reference resources used by multiomeR.** Optional tools are used only when the corresponding input, backend or analysis module is configured. Tools that run before multiomeR are marked as external, and published methods that multiomeR reimplements are marked as multiomeR implementations. General-purpose programming, plotting and workflow-management dependencies are omitted. GEX, gene expression; ATAC, assay for transposase-accessible chromatin; WNN, weighted nearest neighbors; TF, transcription factor.

| Tool/package name [reference] | Purpose in multiomeR |
|---|---|
| Cell Ranger ARC (10x Genomics 2026) | External preprocessing of sequencing reads; supplies aligned reads, GEX counts, ATAC fragments, called barcodes and QC metrics. |
| CellBender (Fleming *et al.* 2023) | Optional external background-RNA removal; corrected GEX matrices are imported in place of Cell Ranger counts. |
| cellsnp-lite (Huang X and Huang 2021) | Optional per-barcode allele counting from the ATAC BAM at donor-VCF variants for genotype demultiplexing. |
| Vireo (Huang Y *et al.* 2019) | Optional assignment of pooled nuclei to donors from allele counts and donor genotypes, including doublet probabilities. |
| BPCells (Parks and Greenleaf 2025) | Disk-backed count and fragment processing, ATAC QC, GEX normalization and PCA, peak counting, ATAC LSI, neighbor graphs and pseudobulk aggregation; optional tile-based peak calling. |
| Seurat / SeuratObject (Hao *et al.* 2024) | Optional SCTransform-based GEX processing and export of multimodal objects containing assays, embeddings, graphs and annotations. |
| sctransform (Hafemeister and Satija 2019) | Variance-stabilizing GEX normalization through the optional Seurat SCTransform backend. |
| Signac (Stuart *et al.* 2021) | ATAC assay and fragment representation in exported objects; its TF–IDF method 1 is implemented on BPCells matrices for LSI. |
| scDblFinder (Germain *et al.* 2022) | Separate GEX- and ATAC-based doublet detection; for ATAC, multiomeR supplies its own peak groups. |
| AMULET (Thibodeau *et al.* 2021) | Fragment-overlap doublet detection using multiomeR's own BPCells-based implementation validated against scDblFinder's AMULET implementation. |
| MACS3 (Zhang Y *et al.* 2008) | Peak discovery from grouped ATAC fragments before the consensus peaks are built. |
| ArchR consensus-peak strategy (Granja *et al.* 2021) | Strategy that multiomeR follows to build summit-centered, fixed-width, non-overlapping consensus peaks for both peak callers. |
| Harmony (Korsunsky *et al.* 2019, Patikas *et al.* 2026) | Optional batch correction of GEX principal-component and ATAC LSI embeddings using configured metadata covariates. |
| Leiden (Traag *et al.* 2019) | Community detection on GEX, ATAC and WNN neighbor graphs, implemented through BPCells. |
| uwot (UMAP) (McInnes *et al.* 2018, Melville 2026) | Low-dimensional visualization of GEX, ATAC and joint WNN representations. |
| Seurat WNN strategy (Hao *et al.* 2021) | Integration of GEX and ATAC neighborhoods with cell-specific modality weights, using multiomeR's own BPCells-based implementation. |
| UCell (Andreatta and Carmona 2021) | Marker-signature scoring for cell-type annotation using multiomeR's own BPCells-based implementation of UCell scoring. |
| JASPAR2026 / motifmatchr (Schep A 2025, Ovek Baydar *et al.* 2026) | Supply motif-family root profiles and identify motif matches in consensus ATAC peaks. |
| betterChromVAR / chromVAR (Schep AN *et al.* 2017, Germain 2026) | Bias-corrected motif-associated accessibility deviations and z-scores for nuclei and pseudobulks; weighted genetic deviations for GWAS annotations. |
| susieR (SuSiE) (Wang *et al.* 2020) | Fine-mapping of candidate regulatory peaks for linked genes, from residualized donor–state pseudobulk accessibility and expression. |
| glmmTMB (Brooks *et al.* 2017) | Donor-level beta-binomial modeling of cell-type composition with configured fixed-effect covariates. |
| edgeR (Robinson *et al.* 2010) | Pseudobulk feature filtering, library-size normalization and quasi-likelihood differential testing of GEX and ATAC counts. |
| limma (Ritchie *et al.* 2015) | Differential modeling of continuous activity scores and blocked count designs, contrast testing and competitive gene-set enrichment with camera. |
| decoupleR / CollecTRI (Badia-i-Mompel *et al.* 2022, Müller-Dott *et al.* 2023) | Inference of TF and TF-complex activity from GEX pseudobulks, with univariate linear models and CollecTRI regulons. |
| msigdbr (Dolgalev 2026) | Retrieval of MSigDB gene sets, including Hallmark (Liberzon *et al.* 2015) and Reactome (Milacic *et al.* 2024) collections, for differential-expression enrichment analysis. |
| Open Targets Platform (Buniello *et al.* 2025) | Retrieval of GWAS credible sets, variant posterior probabilities, study metadata and locus-to-gene evidence for genetic enrichment and interpretation. |
| SCAVENGE (Yu *et al.* 2022) | Trait-relevance scores for each nucleus, from GWAS-associated accessibility spread over the WNN graph, using multiomeR's own sparse-matrix implementation. |

#### Supplementary Table S6

**Resource use of multiomeR and a Seurat/Signac baseline at each aggregation size.** Values behind Supplementary Figure S2, from single runs of each workflow. Supplementary Methods describes the workflows and how each measure was computed. Critical path is estimated from recorded step runtimes, assuming that all of multiomeR's independent steps run at the same time and that the baseline runs its steps one after another. CPU time and RAM are the resources used, sampled from each Slurm job. Peak RAM is the largest memory use of a single step, reached by SCTransform in both workflows. multiomeR's peak at the largest size differed by about 10% between the measured run and an earlier one, more than the workflows differed. multiomeR's values include its multimodal Seurat/Signac export and exclude analyses that the baseline does not perform. Disk includes the Cell Ranger outputs that both workflows read.

| Samples | Input nuclei | Workflow | Critical path (h) | CPU time (CPU-hours) | RAM (GB-hours) | Peak RAM (GB) | Disk (GB) |
|---:|---:|---|---:|---:|---:|---:|---:|
| 1 | 2,713 | multiomeR | 0.04 | 0.15 | 0.30 | 3.4 | 1.2 |
| 1 | 2,713 | Seurat/Signac | 0.08 | 0.14 | 0.16 | 2.6 | 0.7 |
| 2 | 13,609 | multiomeR | 0.15 | 0.85 | 1.70 | 9.1 | 7.7 |
| 2 | 13,609 | Seurat/Signac | 0.41 | 0.83 | 1.64 | 8.2 | 5.3 |
| 5 | 48,922 | multiomeR | 0.37 | 2.50 | 6.51 | 13.0 | 24.8 |
| 5 | 48,922 | Seurat/Signac | 1.39 | 1.84 | 7.12 | 13.9 | 17.4 |
| 10 | 91,191 | multiomeR | 0.56 | 3.75 | 12.24 | 23.4 | 39.3 |
| 10 | 91,191 | Seurat/Signac | 1.79 | 3.74 | 14.56 | 23.4 | 27.9 |
| 20 | 238,222 | multiomeR | 1.17 | 7.18 | 41.20 | 59.2 | 82.3 |
| 20 | 238,222 | Seurat/Signac | 3.57 | 6.28 | 57.91 | 60.6 | 59.6 |
| 31 | 375,757 | multiomeR | 2.17 | 11.31 | 87.25 | 89.0 | 128.4 |
| 31 | 375,757 | Seurat/Signac | 5.57 | 11.34 | 114.24 | 90.4 | 93.4 |
